# Methylenetetrahydrofolate reductase deficiency alters cellular response after ischemic stroke in male mice

**DOI:** 10.1101/857938

**Authors:** Jamie E. Abato, Mahira Moftah, Greg O. Cron, Patrice D. Smith, Nafisa M. Jadavji

## Abstract

**Objective:** Elevated homocysteine concentrations are a risk factor for stroke. A common genetic polymorphism in methylenetetrahydrofolate reductase (*MTHFR* 677 C**➔**T) results in elevated levels of homocysteine. MTHFR plays a critical role in the synthesis of *S*-adenosylmethionine (SAM), a global methyl donor. Our previous work has demonstrated that *Mthfr*^+/−^ mice, which model the *MTHFR* polymorphism in humans, are more vulnerable to ischemic damage. The aim of this study was to investigate the cellular mechanisms by which the MTHFR-deficiency changes the brain in the context of ischemic stroke injury.

**Methods:** In the present study, three-month-old male *Mthfr*^+/−^ and wild-type littermate mice were subjected to photothrombosis (PT) damage. Four weeks after PT damage, animals were tested on behavioral tasks, *in vivo* imaging was performed using T2-weighted MRI, and brain tissue was collected.

**Results:** *Mthfr*^+/−^ animals used their non-impaired forepaw more during to explore the cylinder and had a larger damage volume compared to wild-type littermates. In brain tissue of *Mthfr*^+/−^ mice methionine adenosyltransferase II alpha (MAT2A) protein levels were decreased within the damage hemisphere and increased levels in hypoxia induced factor 1 alpha (HIF-1α) in non-damage hemisphere. There was an increased antioxidant response in the damage site as indicated by higher levels of nuclear factor erythroid 2-related factor 2 (Nrf2) and superoxide dismutase 2 (SOD2).

**Conclusions:** Our results suggest that *Mthfr*^+/−^ mice are more vulnerable to PT-induced stroke damage through regulation of the cellular response. The increased antioxidant response we observed may be compensatory to the damage amount.

## 1. Introduction

Stroke is a leading cause of death worldwide (Durga et al., 2007) and predominantly targets older adults (Katan and Luft, 2018). As the population worldwide ages, the prevalence of stroke is expected to increase (Mozaffarian et al., 2015; Katan and Luft, 2018). Older adults are at a higher risk of stroke, and also have a reduced ability to absorb all the nutrients from their diet (Bottiglieri et al., 2000; Shlisky et al., 2017).

Reduced levels of B-vitamins, a major component of one-carbon metabolism, result in elevated homocysteine concentrations which are associated with increased risk for cardiovascular disease, such as stroke (Castro et al., 2006; Lehotský et al., 2016). B-vitamins, include folic acid and vitamin B12, play an important role in reducing levels of homocysteine, through one-carbon metabolism. Metheylenetetrahydrofolate reductase (MTHFR) is an enzyme that catalyzes the irreversible conversion of 5,10-methylenetetrahydrofolate to 5-methyltetrahydrofolate (5-methylTHF). The methyl group from 5-methylTHF is a substrate in the vitamin-B12-dependent methylation of homocysteine to form methionine by methionine synthase. A polymorphism in *MTHFR* (677 C**➔**T) (Frosst et al., 1995), which is present in 5-20% of North American and European populations (Schneider et al., 1998), results in elevated levels of homocysteine. The polymorphism has been associated with increased risk for vascular disease (Frosst et al., 1995) including ischemic stroke (Casas et al., 2005; Cronin et al., 2005; Song et al., 2016; Azad et al., 2017; Li et al., 2017; Mao and Han, 2018). However, the mechanism through which this increased vulnerability exists is not well defined.

The *Mthfr*^+/−^ mouse models traits of the human polymorphism, including reduced levels of enzyme activity and elevated levels of homocysteine (Chen et al., 2001; Jadavji et al., 2012). Using the MTHFR-deficient mice and the photothrombosis induced ischemic damage, our previous research has demonstrated that aged *Mthfr*^+/−^ mice are more vulnerable to damage after ischemic stroke through reduced neuronal and astrocyte viability (Jadavji et al., 2017, 2018, 2019). The mechanisms through which the *Mthfr*^+/−^ mice are more impaired after ischemic damage remains uncharacterized. The aim of this study was to investigate the cellular response through which MTHFR deficiency regulates the brain’s vulnerability to ischemic stroke.

## 2. Experimental Design

### 2.1 Animals and experimental design

All experiments were conducted in accordance with the guidelines of the Canadian Council on Animal Care (CCAC). Two cohorts of male C57Bl/6 *Mthfr* heterozygote (*Mthfr*^+/−^, n = 15) and wild-type littermate controls (*Mthfr*^+/+^, n = 15) mice were housed in standard caging conditions with *ad libitum* standard mouse chow (Envigo) and water for the duration of the experiment. Animals were randomly assigned to each of the experimental groups.

### 2.2 Photothrombosis

At 3-months of age, animals were subjected photothrombosis (PT) induced ischemic damage. Focal ischemic damage was targeted to the sensorimotor cortex (mediolateral ± 0.24 mm) as previously described (Lee et al., 2004; Jadavji et al., 2017, 2018). Animals were anaesthetized with isoflurane (1.5%) in a 70:30 nitrous oxide: oxygen mixture. Core body temperature was maintained at 37.0 ± 0.2°C using an automated heat blanket with temperature feedback (Harvard Apparatus). An intraperitoneal injection of 10 mg/kg photoactive Rose Bengal (Sigma) was administered 5 min prior to a 532 nm green laser (Beta Electronics) exposure on the skull.

### 2.3 Behavioral Testing

#### 2.3.1 Bederson exam

The Bederson exam evaluates symmetry, forepaw outstretching, climbing, and reflex (Bederson et al., 1986). A perfect score equates to a zero. Neurological testing was performed 4 weeks after PT damage and investigators were blinded to the experimental groups.

#### 2.3.2 Forepaw Asymmetry

A cylinder (19 cm high, 14 cm in diameter) made of thick glass was used to assess forepaw asymmetry. The mouse was placed in the glass cylinder for 10 min and video was recorded from the side. Forepaw contacts with the wall of the cylinder were counted. The final score was calculated as follows: final score = (number of non-impaired forelimb movement – number of impaired forelimb movement)/ (sum of all movements). A positive score thus indicated favored use of the non-impaired forelimb, a negative indicated favored use of the impaired forelimb and a score of zero indicated equal use of both non-impaired and impaired forelimbs upon rearing and exploration of the cylinder walls (Theoret et al., 2015).

### 2.4 MRI lesion volume analysis

At four weeks after PT damage, lesion volume was quantified. A 7 Tesla GE/Agilent MR 901 was used to collect the data. Mice were anaesthetized prior to the MRI procedure using isoflurane. A 2D fast spin echo sequence (FSE) pulse sequence was used for the imaging, with the following parameters: slice thickness = 0.5 mm, spacing = 0 mm, field of view = 2.5 cm, matrix = 256 × 256, echo time = 41 ms, repetition time = 4000 ms, echo train length = 8, bandwidth = 16 kHz, 2 averages, fat saturation, imaging time = 5 minutes.

### 2.5 Western blot

At the completion of experiments, approximately 4 weeks after PT damage, animals were decapitated to obtain fresh brain tissue collection for Western blot analysis, PT and non-PT regions (~50 mg) were micro-dissected. RIPA buffer containing phosphatase and proteinase inhibitors (Cell Signalling) was used to extract proteins. Proteins were separated by SDS-PAGE and transferred to nitrocellulose membranes. Primary antibodies were; anti-rabbit methionine adenosyl transferase (MAT2A) (1:2000; AbCam), anti-rabbit hypoxia induced factor 1 alpha (HIF-1α) (1:500; AbCam), or anti-rabbit glyceraldehyde 3-phosphate dehydrogenase (GAPDH) (1:10 000; Cell Signalling) overnight. Primary antibodies were visualized with infrared dye goat anti-rabbit IgG (1:10 000, 800 Channel; LiCor Biosciences). Imaging was performed using Near Infrared (NIR) detection (Li-Cor Biosciences). Quantification of bands was determined using densitometry analysis via ImageJ (National Institutes of Health) and bands were normalized to GAPDH.

### 2.6 Immunofluorescence

After the completion of behavioral testing, animals were euthanized using the perfusion method. Brain tissue was fixed in 4% paraformaldehyde (PFA) for 24 hrs. Using a cryostat, brain tissue was sectioned at 30 μm thickness then mounted on slides in serial order.

To explore mechanisms of neurodegeneration, levels of oxidative stress using nuclear factor erythroid 2-related factor 2 (NRF2) and superoxide dismutase 2 (SOD2) were investigated. Primary antibodies used were rabbit anti-NRF2 (1:100, AbCam), rabbit anti-SOD2 (1:100, Life Technologies), and mouse anti-neuronal nuclei (NeuN) (1:200, AbCam) diluted in 0.5% Triton X. Brain tissue was incubated overnight at 4°C with primary antibodies. The following day, secondary antibodies (1:200, Cell Signaling Technologies) brain tissue was incubated at room temperature for 2 h and then stained with 4’, 6-diamidino-2-phenylindole (DAPI) (1:10, 000). For analysis in within the ischemic core of brain tissue, co-localization of NRF2 or SOD2 with NeuN labelled neurons were counted and averaged per animal. Data from five animals per group were analyzed.

### 2.7 Statistics analysis

All data analysis was conducted by two individuals blinded to experimental groups. Data from the study were analyzed using GraphPad Prism 6.0. Unpaired t-tests were performed on behavioral testing, MRI, immunofluorescence, and Western Blot experiments between genotype groups with PT damage. In all analysis, a p-value less than or equal to 0.05 was considered significant. All data are presented as mean ± standard error of the mean (SEM).

## 3. Results

### 3.1 Forepaw usage impairs in Mthfr^+/−^ mice

There was no difference in the neurological testing score between *Mthfr*^+/+^ and *Mthfr*^+/−^ mice (Figure 1A, t (29) = 1.62, p = 0.1). However, *Mthfr*^+/−^ mice showed a preference to use their non-impaired forelimb, while completing the forepaw asymmetry task compared to *Mthfr*^+/+^ animals (Figure 1B, t (29) = 3.46, p < 0.01).

**Figure 1.**
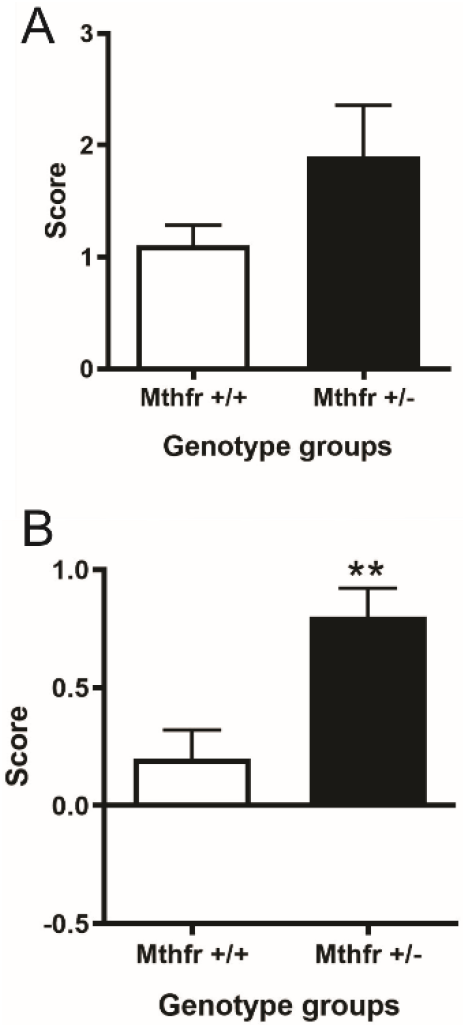
Behavioral function in adult *Mthfr*^+/+^ and *Mthfr*^+/−^ animals 4 weeks after PT damage. Neurological testing (A) and forepaw asymmetry (B) scores in *Mthfr*^+/+^ and *Mthfr*^+/−^ mice. Bars represent mean ± SEM of 15 animals per group. ** p < 0.01 un-paired t-test between *Mthfr*^+/+^ and *Mthfr*^+/−^ mice.

**Figure 2.**
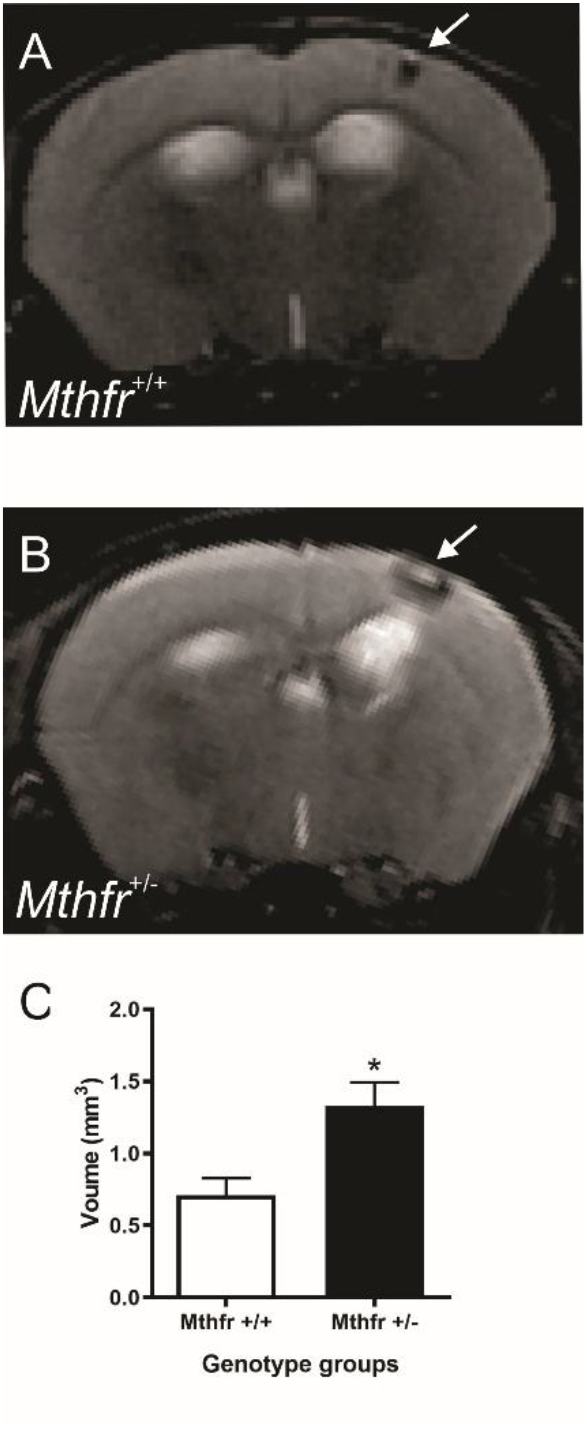
The impact of MTHFR-deficiency on damage volume 4-weeks after PT damage to the sensorimotor cortex. Representative MRI T2-weighted images from *Mthfr*^+/+^ (A) and *Mthfr*^+/−^ (B) mice. Quantification of lesion volume (C). Bars represent mean ± SEM of 4 animals per group. * p < 0.05 un-paired t-test between *Mthfr*^+/+^ and *Mthfr*^+/−^ mice.

### 3.2 Increased damage volume in Mthfr^+/−^ mice

Quantification of PT lesion volume using T2 weighted MRI revealed that the *Mthfr*^+/−^ mice showed a larger lesion volume in the sensorimotor cortex compared to wildtype animals (Figure 1C, t (7) = 3.1, p<0.05).

### 3.3 Altered cellular response within photothrombosis damage area of Mthfr^+/−^ mice

We examined two markers of cellular response after PT damage, the first an enzyme, MAT2A, which is linked to MTHFR through one-carbon metabolism and is important in the generation of *S*-adenosylmethionine (SAM), a global methyl donor. Four weeks after PT damage we measured levels of MAT2A. In *Mthfr*^+/−^ mice there was a decrease in levels of MAT2A within the PT damage hemisphere compared to *Mthfr*^+/+^ animals (Figure 3A, t (13) = 2.7, p < 0.05). There were no changes in MAT2A within the non-PT hemisphere between groups (data not shown, t (13) =11.4, p = 0.58).

**Figure 3.**
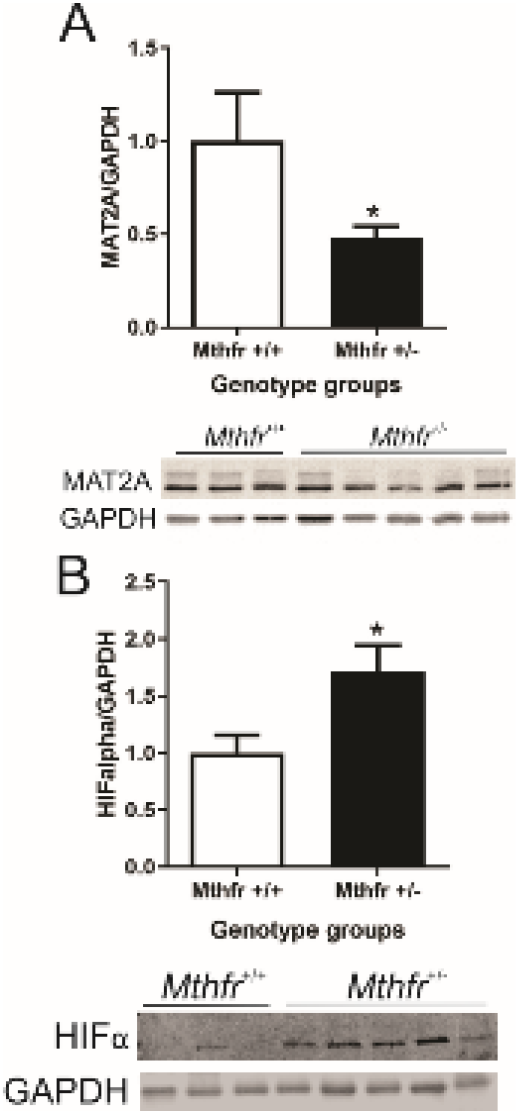
The impact of MTHFR-deficiency after ischemic damage on cellular responses to ischemia. Protein levels of methionine adenosyltransferase 2A (MAT2A) in PT hemisphere (A), and non-PT hemisphere levels of hypoxia-inducible factor 1-alpha (HIF-1α) (B) in *Mthfr*^+/+^ and *Mthfr*^+/−^ animals 4-weeks after PT damage. Bars represent mean ± SEM of 7 mice per group. The panels below the graphs depict a representative Western Blot. *p<0.05 and **p<0.01 indicate Tukey’s pairwise comparison between indicated groups.

HIF-1α is an regulator of the cellular response after hypoxia (Katsimpardi et al., 2014). Previously we have shown increased levels of HIF-1α *Mthfr*^+/−^ and *Mthfr*^−/−^ primary astrocytes after exposure to hypoxia (Jadavji et al., 2018), as well as in brain tissue of *Mthfr*^+/−^ mice with ischemic stroke (Jadavji et al., 2019). In the present study, using an *in vivo* model of MTHFR deficiency and ischemic damage, we observed increased levels of HIF-1α within the non-damage PT hemisphere of *Mthfr*^+/−^ mice (Figure 3B, t (13) = 3.4, p < 0.01). Interestingly, we did not observe any expression of HIF-1α within the lesion hemisphere of *Mthfr*^+/+^ and *Mthfr*^+/−^ animals (data not shown).

### 3.4 Increased oxidative stress within damage area of Mthfr^+/−^ mice

We measured oxidative stress 4 weeks after damage in sectioned brain tissue of animals, by assessing levels of nuclear factor (erythroid-derived 2)-like 2 (Nrf2) and superoxide dismutase 2 (SOD2). Analysis of oxidative stress was conducted within the damage area. Representative images of Nrf2 are shown in Figure 4A and B. There were increased levels of Nrf2 within the damaged area of *Mthfr*^+/−^ mice (Figure 4C, t (9) = 4.02, p < 0.001). Representative SOD2 images are shown in figures 4D and 4E. *Mthfr*^+/−^ mice had increased levels of SOD2 in brain tissue (Figure 4F, t (9) = 2.57, p < 0.05).

**Figure 4.**
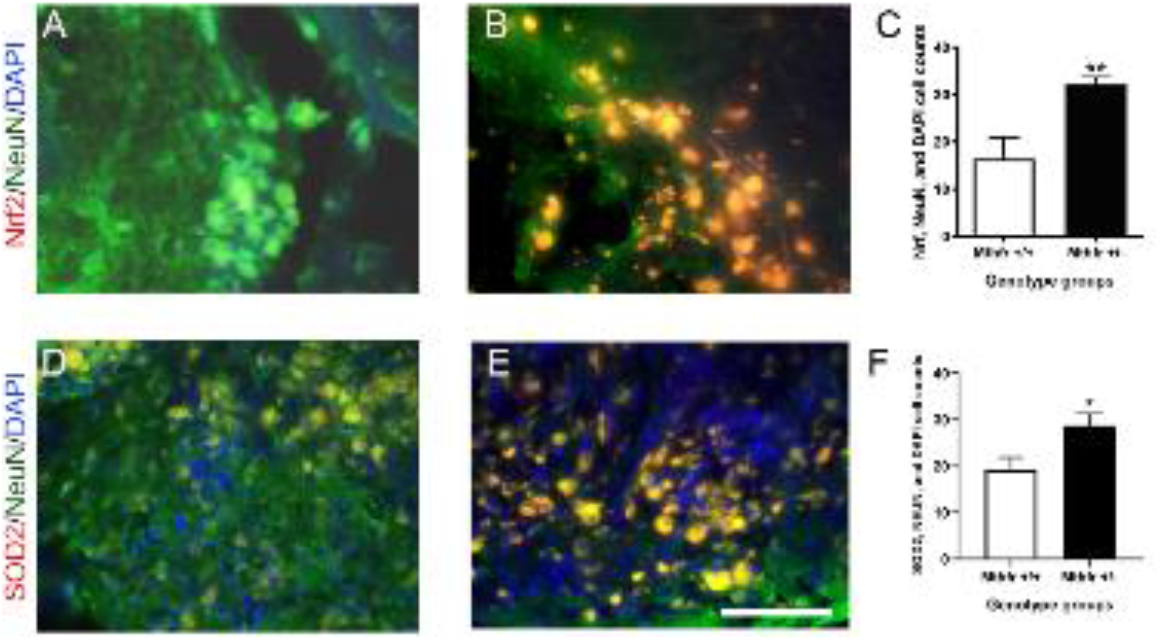
Antioxidant activity within ischemic area of MTHFR-deficient mice. Representative images of nuclear factor (erythroid-derived 2)-like 2 (Nrf2), (A) and (B). Quantification of Nrf2, neuronal nuclei (NeuN), and DAPI (4′,6-diamidino-2-phenylindole) positive cells (C). Representative superoxide dismutase 2 (SOD2) images, (D) and (E). Quantification of SOD2, NeuN and DAPI positive cells within damage area (F). Bars represent mean ± SEM of 5 mice per group. *p<0.05 and **p<0.01 indicate unpaired t-test comparison between indicated groups.

## 4. Discussion

Elevated levels of homocysteine are a risk factor for stroke (Castro et al., 2006). Previously we have reported that aged *Mthfr*^+/−^ mice are more vulnerable to ischemic damage (Jadavji et al., 2018). We have also shown this increased vulnerability *in vitro* using primary neurons (Jadavji et al., 2017) and astrocytes (Jadavji et al., 2018). In the present study, we have confirmed our previous findings that *Mthfr*^+/−^ mice have a more severe motor impairment after ischemic stroke using the forepaw asymmetry task. The heterozygote mice also had increased damage volume within the sensorimotor cortex. To understand the cellular response after ischemic damage, we report reduced levels of MAT2A *Mthfr*^+/−^ mice within the damage area. MAT2A is an enzyme linked to MTHFR through one-carbon metabolism and involved in generating SAM within the damage brain region. Within the non-lesion hemisphere there are increased levels of HIF-1α in the non-damaged PT area. HIF-1α is involved in cellular responses for protection and repair. The anti-oxidative response was increased in within the damage of *Mthfr*^+/−^ mice.

In patients with vascular disease, changes in DNA methylation have been reported (Castro et al., 2003). Using animal models to understand diseases processes, it has been suggested that decreases in methylation may promote cell death after stroke (Kalani et al., 2013). Furthermore, changes in methylation may impact stroke onset and progression (Endres et al., 2000; Kalani et al., 2013). MTHFR generates methyl groups, and the impact of a MTHFR-deficiency on methylation after stroke remains unclear; our present study has attempted to fill this gap. Previous work reported that *Mthfr*^+/−^ mice have increased levels of hypomethylation in the brain (Chen et al., 2001). MAT2A is a catalytic subunit of methionine adenosyltransferase (MAT), an essential enzyme for the catalysis of the principle biological methyl donor, SAM from methionine and ATP. In the present study, we observed increased levels of MAT2A in wild-type mice which is in line with increased methylation after ischemic stroke that has previously been reported (Endres et al., 2000; Kalani et al., 2013). Interestingly, lower levels of MAT2A may lead to reduced levels of SAM, the global methyl donor (Frau et al., 2012), however this requires further confirmation.

HIF-1α is a master regulator of cell response to hypoxia, since it leads to the expression of several genes involved in the adaptation of oxygen availability and angiogenesis (Jiang et al., 2001; Cunningham et al., 2012). In the kidney, HIF-1α stabilizes cells after ischemia and interference with HIF-1α promotes tissue damage and dysfunction (Conde et al., 2012). In the present study, we observed increased levels of HIF-1α within the non-PT hemisphere in *Mthfr*^+/−^ mice. These results support our previous work, in which we report increased HIF-1α levels in primary astrocyte cultures from *Mthfr*^+/−^ and *Mthfr*^−/−^ embryos after a hypoxia treatment (Jadavji et al., 2018). Increased levels of HIF-1α in the non-PT hemisphere may be another adaptive response to compensate for damage in the other hemisphere.

In the brain, both ischemic stroke (Siti et al., 2015) and increased levels of homocysteine (Hoffman, 2011) result in oxidative stress. The anti-oxidative response is important in regulating the impact of oxidative stress in brain tissue especially after damage, during neurodegeneration (Zhang and Butterfield, 2017). Our previous work has shown that dietary supplementation of one-carbon metabolites including folic acid (vitamin B9), choline, vitamin B12, and riboflavin (vitamin B2) increase the anti-oxidative response in brain tissue after ischemic stroke (Jadavji et al., 2017). Targeting anti-oxidants after stroke in patients is complicated (Siti et al., 2015), but increasing activation of Nrf2 could be a potential target (Zhang et al., 2017). Additionally, the combination of anti-oxidant through diet and pharmaceutical therapies may be beneficial for stroke-affected patients.

Together, the data from this study provides insight on potential mechanisms that enhance stroke vulnerability in *Mthfr*^+/−^ mice. Ischemic stroke in MTHFR-deficient animals changes cellular responses in brain tissue, making *Mthfr*^+/−^ mice more vulnerable to damage. Further investigation is required to dissect the impact of changes to SAM levels and consequential global genetic silencing processes in MTHFR-deficiency that contribute to ischemic stroke outcomes.

## Acknowledgements

This research was supported by Natural Sciences and Engineering Research Council of Canada (NMJ and PDS) and Midwestern University start-up funds (NMJ).

## Disclosure statement

The authors report no conflict of interest

## Notes

#### Summary of Updates

Revisions to manuscript text.

